# Profiling the expression and function of ER46 in human endometrial tissues and uterine NK cells

**DOI:** 10.1101/777607

**Authors:** Douglas A Gibson, Arantza Esnal-Zufiaurre, Cristina Bajo-Santos, Frances Collins, Hilary OD Critchley, Philippa TK Saunders

## Abstract

**Study question:** Does the oestrogen receptor isoform, ER46, contribute to regulation of endometrial function?

**Summary answer:** ER46 is expressed in endometrial tissues during the proliferative and secretory phases and is the predominant ERα isoform in first trimester decidua. ER46 is abundantly expressed in uterine NK (uNK) cells and localised to the cell membrane. Activation of ER46 regulates the function of human uNK cells by increasing cell motility.

**What is known already:** Oestrogens acting via their cognate receptors are essential regulators of endometrial function and play key roles in establishment of pregnancy. ER46 is a 46kDa truncated isoform of full length ERα (ER66, encoded by *ESR1*) that contains both ligand and DNA binding domains. Expression of ER46 in human endometrium has not been investigated previously. ER46 is located at the cell membrane of peripheral blood leukocytes and mediates rapid responses to oestrogens. UNK cells are a phenotypically distinct (CD56^bright^CD16^-^) population of tissue-resident immune cells that regulate vascular remodelling within the endometrium and decidua. We have shown that oestrogens stimulate rapid increases in uNK cell motility. Previous characterisation of uNK cells suggests they are ER66-negative but expression of ER46 has not been characterised. We hypothesise that uNK cells express ER46 and that rapid responses to oestrogens are mediated via this receptor.

**Study design, size, duration:** This laboratory-based study used primary human endometrial (n=24) and decidual tissue biopsies (n=30) as well as uNK cells which were freshly isolated from first trimester human decidua (n=18).

**Participants/materials, setting, methods:** Primary human endometrial and first trimester decidual tissue biopsies were collected using methods approved by the local institutional ethics committee (LREC/05/51104/12 and LREC/10/51402/59). The expression of oestrogen receptors (ER66, ER46 and ERβ) was assessed by qPCR, western blot and immunohistochemistry. Uterine Natural Killer (uNK) cells were isolated from first trimester human decidua by magnetic bead sorting. Cell motility of uNK cells was measured by live cell imaging: cells were treated with oestradiol (E2)-BSA (10nM equivalent), the ERβ-selective agonist 2,3-bis (4-hydroxyphenyl)-propionitrile (DPN; 10nM) or vehicle control (DMSO).

**Main results and the role of chance:** ER46 was detected in proliferative and secretory phase tissues and was the predominant ERα isoform in first trimester decidua samples. Immunohistochemistry revealed ER46 was co-localised with ER66 in cell nuclei during the proliferative phase but detected in both the cytoplasm and cell membrane of stromal cells in the secretory phase and in decidua. Triple immunofluorescence staining of decidua tissues identified expression of ER46 in the cell membrane of CD56-positive uNK cells which were otherwise ER66-negative. Profiling of isolated uNK cells confirmed expression ER46 and localised ER46 protein to the cell membrane. Functional analysis of isolated uNK cells using live cell imaging demonstrated that activation of ER46 with E2-BSA significantly increased uNK cell motility.

**Limitations, reasons for caution:** Expression patterns in endometrial tissue was only determined using samples from proliferative and secretory phases. Assessment of first trimester decidua samples was from a range of gestational ages which may have precluded insights into gestation specific changes in these tissues. Our results are based on *in vitro* responses of primary human cells and we cannot be certain that similar mechanisms occur *in situ*.

**Wider implications of the findings:** E2 is an essential regulator of reproductive competence. This study provides the first evidence for expression of ER46 in human endometrium and decidua of early pregnancy. We describe a mechanism for regulating the function of human uNK cells via expression of ER46 and demonstrate that selective targeting with E2-BSA regulates uNK cell motility. These novel findings identify a role for ER46 in human endometrium and provide unique insight into the importance of membrane-initiated signalling in modulating the impact of E2 on uNK cell function in women.

**Study funding/competing interest(s):** These studies were supported by MRC Programme Grants G1100356/1 and MR/N024524/1 to PTKS. HODC was supported by MRC grant G1002033.

## Introduction

Oestrogens are essential for reproductive function and fertility. They classically mediate their functions by binding to cognate receptors, ERα and ERβ, encoded by the genes *ESR1* and *ESR2* respectively. Oestrogens act via systemic endocrine signals and via local intracrine action to regulate key functional processes within the endometrium including proliferation, angiogenesis and inflammation (Gibson, et al., 2012) that prime the endometrium for establishment and maintenance of pregnancy (Gibson, et al., 2013, Gibson, et al., 2018). Oestrogen action is controlled by ligand availability but also via expression and localisation of ER isoforms which are altered in a cell and tissue context-dependent manner. We have previously used qPCR and immunohistochemistry to document stage and cell-specific expression of ERα and ERβ, as well as ERβ splice variant isoforms in human endometrium and decidua of early pregnancy (reviewed in (Gibson, et al., 2012)). Endometrial ERα expression is greatest in the proliferative phase with decreased expression in the secretory phase and a further reduction in first trimester decidual tissue compared to non-pregnant endometrial tissues (Critchley, et al., 2002, Milne, et al., 2005). In those studies we used a mouse monoclonal antibody directed against recombinant human ERα; the epitope for this antibody was not defined but it recognised a protein of 66KDa (consistent with full length wild type ERα) in breast cancer cell homogenates detected by western blot (Chantalat, et al., 2016) and detected ERα in both stromal and epithelial cells by immunohistochemistry ((Bombail, et al., 2008) see figures 4 and 5). In these studies, immunostaining for ERα detected a protein that was exclusively nuclear, consistent with the established functional role of this receptor protein as a ligand-activated transcription factor.

**Figure 1.**
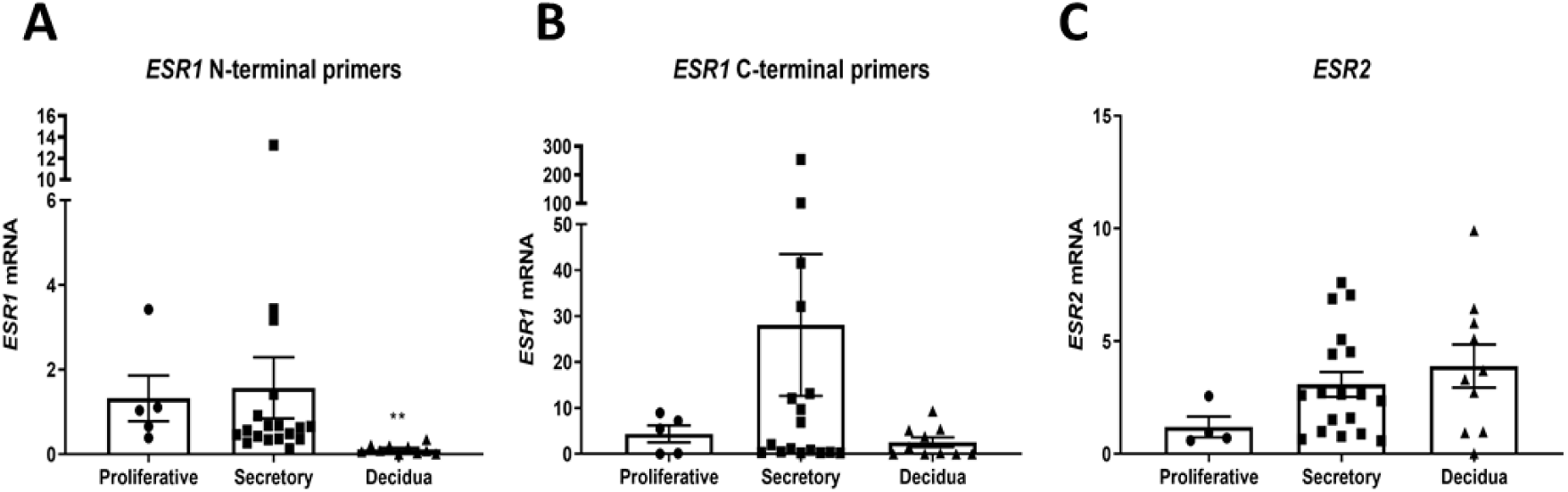
Expression of ER isoforms in endometrial tissues. The expression of *ESR1*, using N- and C-terminal primers, and *ESR2* was assessed using qPCR in proliferative and secretory phase endometrium as well as first trimester decidua tissue samples. **A** N-terminal primers detected mRNAs encoding *ESR1* in all endometrial tissues, expression was unchanged between proliferative and secretory endometrial tissues and significantly decreased in decidua. **B** C-terminal primers detected mRNAs encoding *ESR1* in all endometrial tissues, expression was unchanged between endometrial tissues but mean expression of *ESR1* was greatest in secretory phase endometrial samples. **C** *ESR2* was detected in all endometrial tissues. Tissues for qPCR analysis; proliferative, *n* = 4-5; secretory *n* = 18; decidua, *n* = 10. Kruskal–Wallis test with multiple comparisons. \*\**P* < 0.01. All data are presented as mean ± s.e.m.

**Figure 2.**
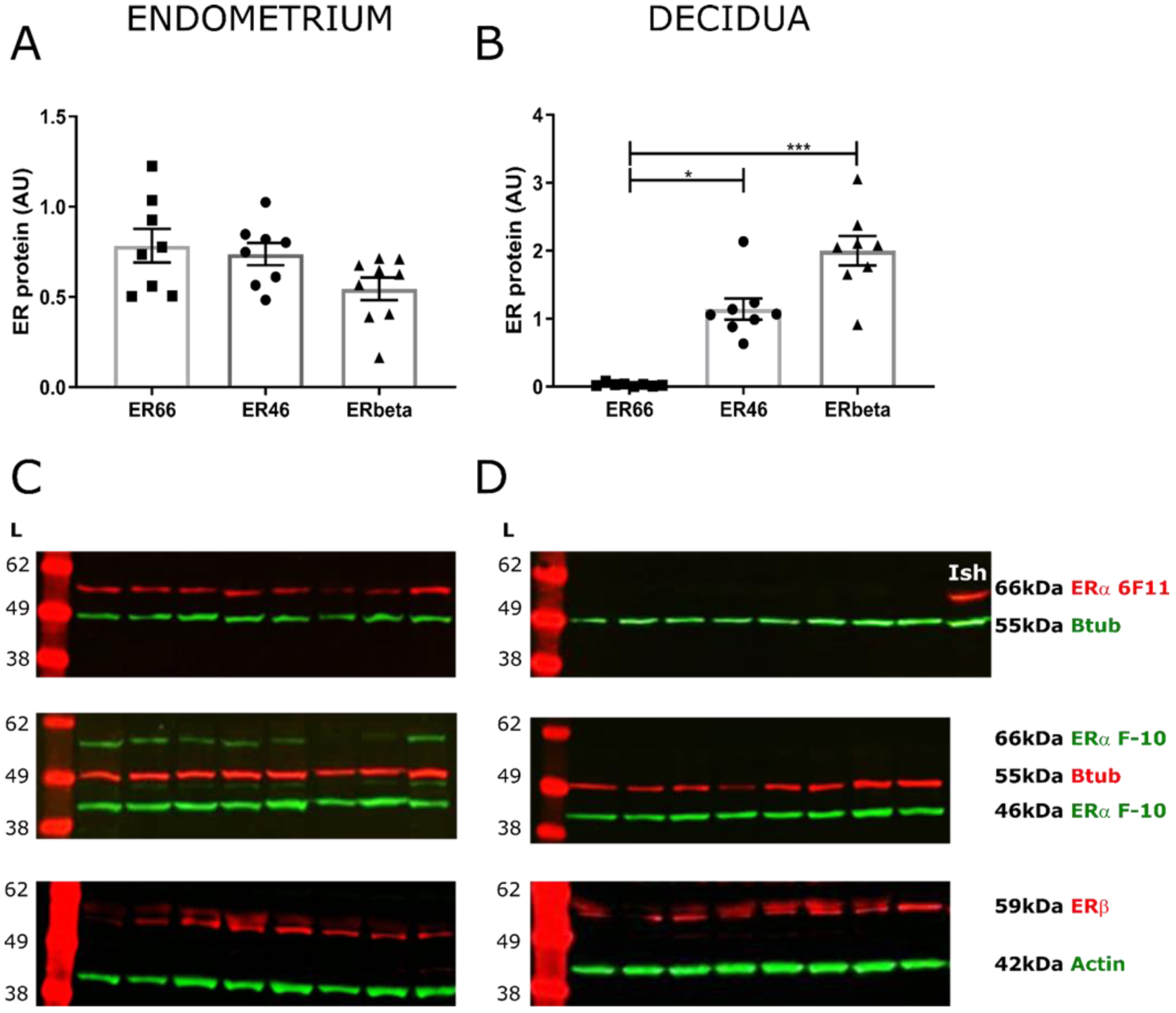
Protein expression of ER isoforms in endometrial tissues. Protein expression of ER isoforms in endometrial tissue homogenates (proliferative phase n=4, secretory phase n=4, decidua n=8) was assessed by western blot using the ERα 6F11 or the C-terminal-specific ERα F-10 antibodies and an antibody that detected ERβ. **A** Protein expression of ER isoforms ER66 (6F11 antibody band), ER46 (F-10 antibody band) and ERβ was assessed by densitometry analysis in endometrium (**A**) and decidua (**B**) and normalised to loading control. All isoforms were present in endometrial tissues (pooled proliferative and secretory) but ER46 (p<0.05) and ERβ (p<0.001) protein concentrations were significantly greater than ER66 in decidual tissues. **C** In non-pregnant endometrial tissues, full length ERα was detected at a band corresponding to 66kDa (red) with the ERα 6F11 antibody. The C-terminal ERα F-10 antibody detected two bands in non-pregnant endometrial tissue homogenates corresponding to 66kDa and 46kDa (green). ERβ was detected at a band corresponding to 59 kDa (red). **D** In first trimester decidua tissue homogenates, full length ERα was not detected with the ERα 6F11 antibody but was present in Ishikawa cell control homogenate (Ish) at a band corresponding to 66kDa (red). Only the 46kDa band was detected in decidua tissue homogenates using the ERα F-10 antibody (green). ERβ was detected at a band corresponding to 59 kDa (red). ER antibodies, band sizes and loading controls (actin or B-tubulin (Btub)) as indicated. AU - arbitrary units. \**P* < 0.05. \*\*\**P* < 0.001. Kruskal–Wallis test with multiple comparisons. Data are presented as mean ± s.e.m.

**Figure 3.**
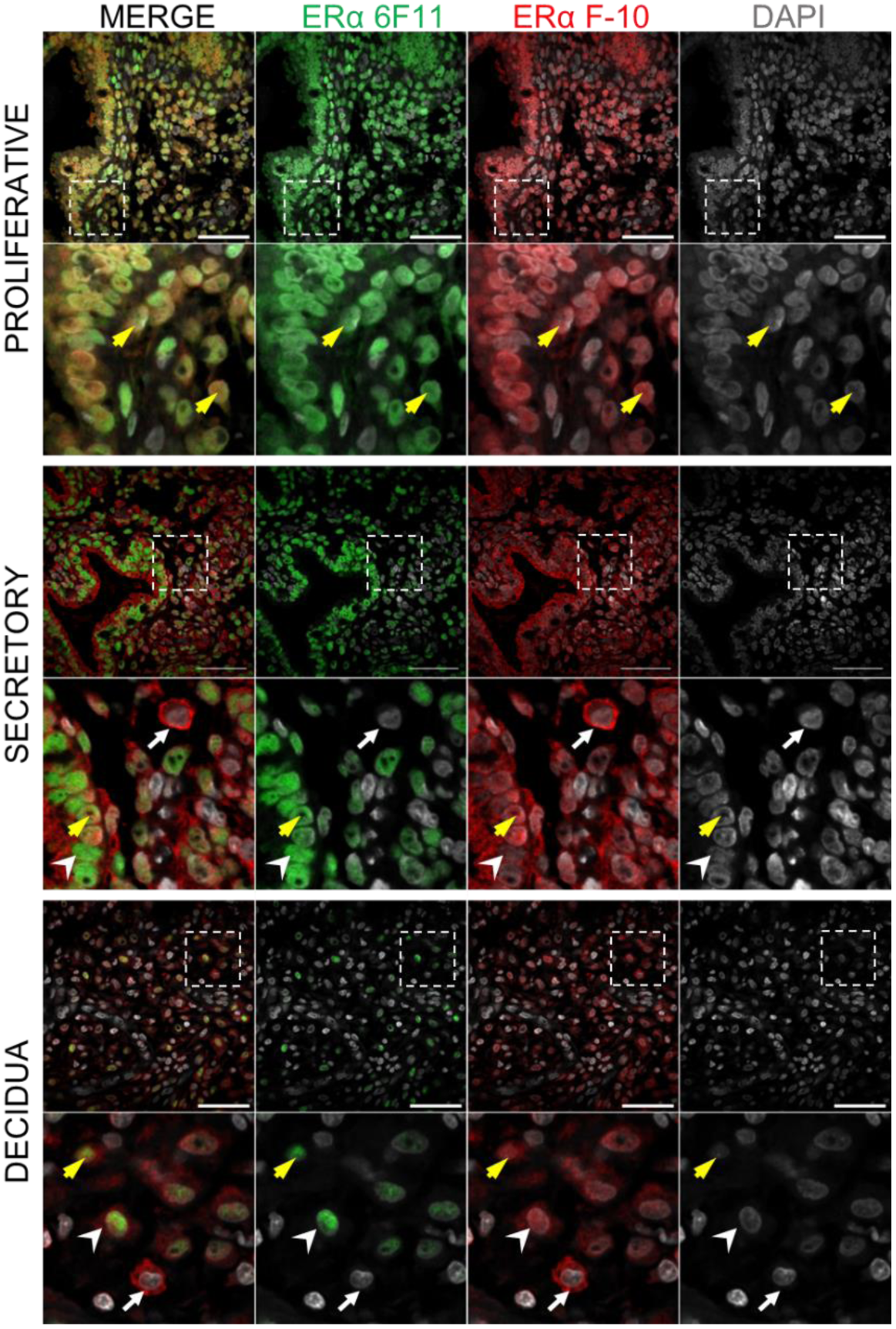
Immunostaining of ER isoforms in endometrial tissues. The expression and localisation of ER proteins in endometrial tissues was assessed using multiplex immunohistochemistry. The ERα 6F11 (green) or the C-terminal-specific ERα F-10 antibodies (red) identified either ER66 or both ER66 and ER46 respectively. In proliferative phase endometrial biopsies, the expression of both antibodies co-localised and was detected in the nuclei of all cells (inset; yellow arrows). In secretory phase endometrial biopsies, strong nuclear staining for ER66 was detected using the ERα 6F11 antibody in both stromal and epithelial cells (arrowhead; green) and co-localisation of both antibodies was detected in the nuclei of some stromal cells (yellow arrow). Extra-nuclear expression of ERα (putatively ER46) was detected in the cytoplasm of epithelial and stromal cells (red) and was localised to the membrane of some cells within the stromal compartment which did not express ER66 (putative immune cells; white arrow). In decidua tissues, extra-nuclear expression of ERα (F-10 ERα antibody; putatively ER46) was detected in the cytoplasm of stromal cells (red; arrowhead) and was localised to the membrane of putative immune cells which did not express ER66 (white arrow). Some nuclear expression of ER66 was detected using by the ERα 6F11 antibody in stromal cells (green) and co-expression of both antibodies was detected in the nuclei of stromal cells (yellow arrow). Dashed box indicates cropped zoom region. Images are representative of at least 3 different patient samples per tissue type. Scale bars 20 µm, nuclear counterstain DAPI (grey).

**Figure 4.**
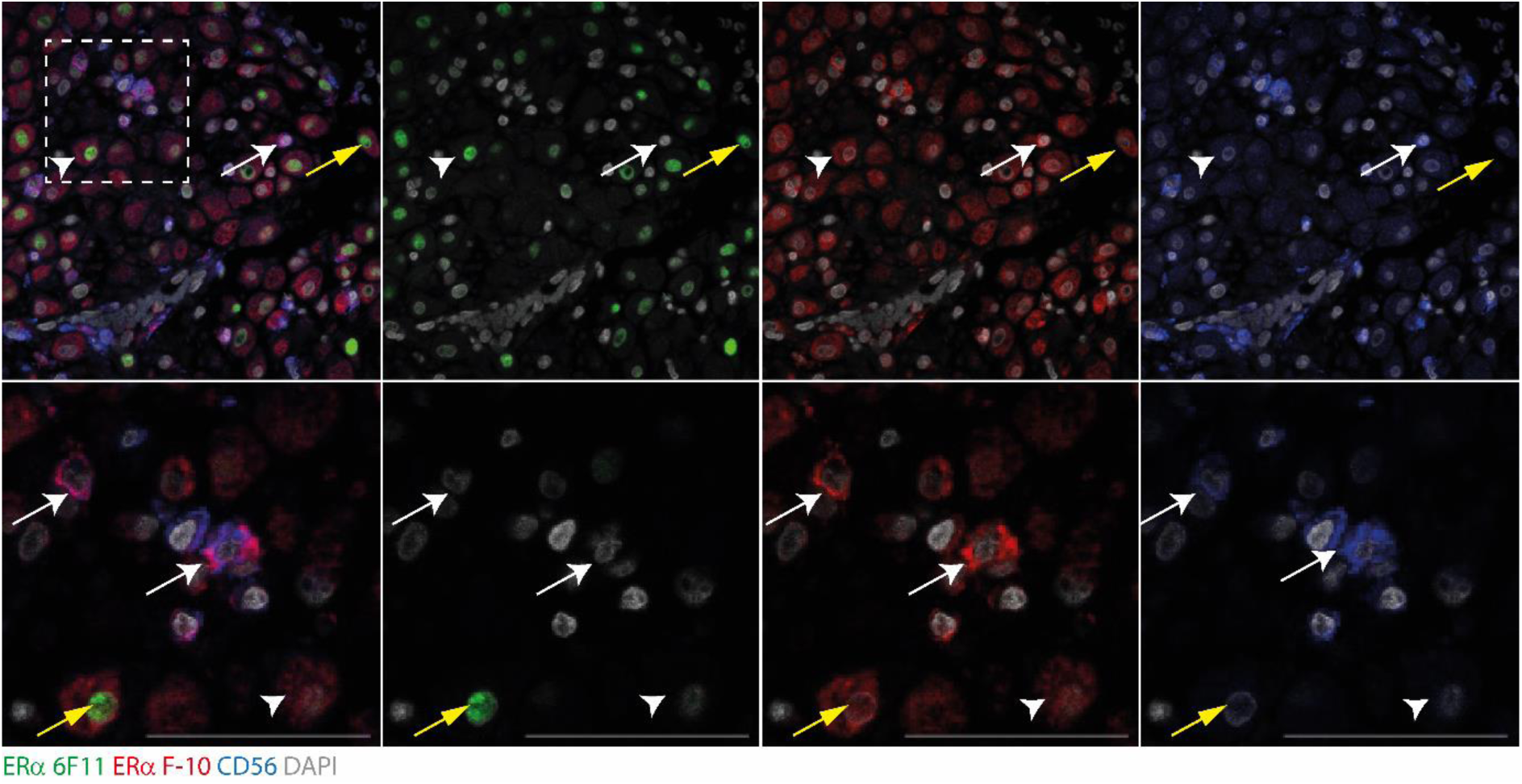
Expression of ER46 in decidual uNK cells. The expression and localisation of ER proteins in uNK cells was assessed in decidua tissues using multiplex immunohistochemistry. The ERα 6F11 (green) or the C-terminal-specific ERα F-10 antibodies (red) and the uNK cell marker CD56 (blue) were assessed. UNK cells were abundant in decidua and staining for surface marker CD56 (blue) co-localised with membrane staining for ERα (red) identified using the ERα F-10 antibody (ER46; white arrows) but were negative for ER66. ERα identified using the ERα F-10 antibody was also detected in the cytoplasm of stromal cells and weakly in stromal nuclei (red, white arrowhead). Some nuclear staining for ER66 was detected using the ERα 6F11 antibody in stromal cells (green) which co-expressed ER46 detected with ERα F-10 antibody staining (yellow arrow). Images are representative of staining from at least 3 different patient samples. Dashed box indicates cropped zoom region. Scale bars 50 µm, nuclear counterstain DAPI (grey).

**Figure 5.**
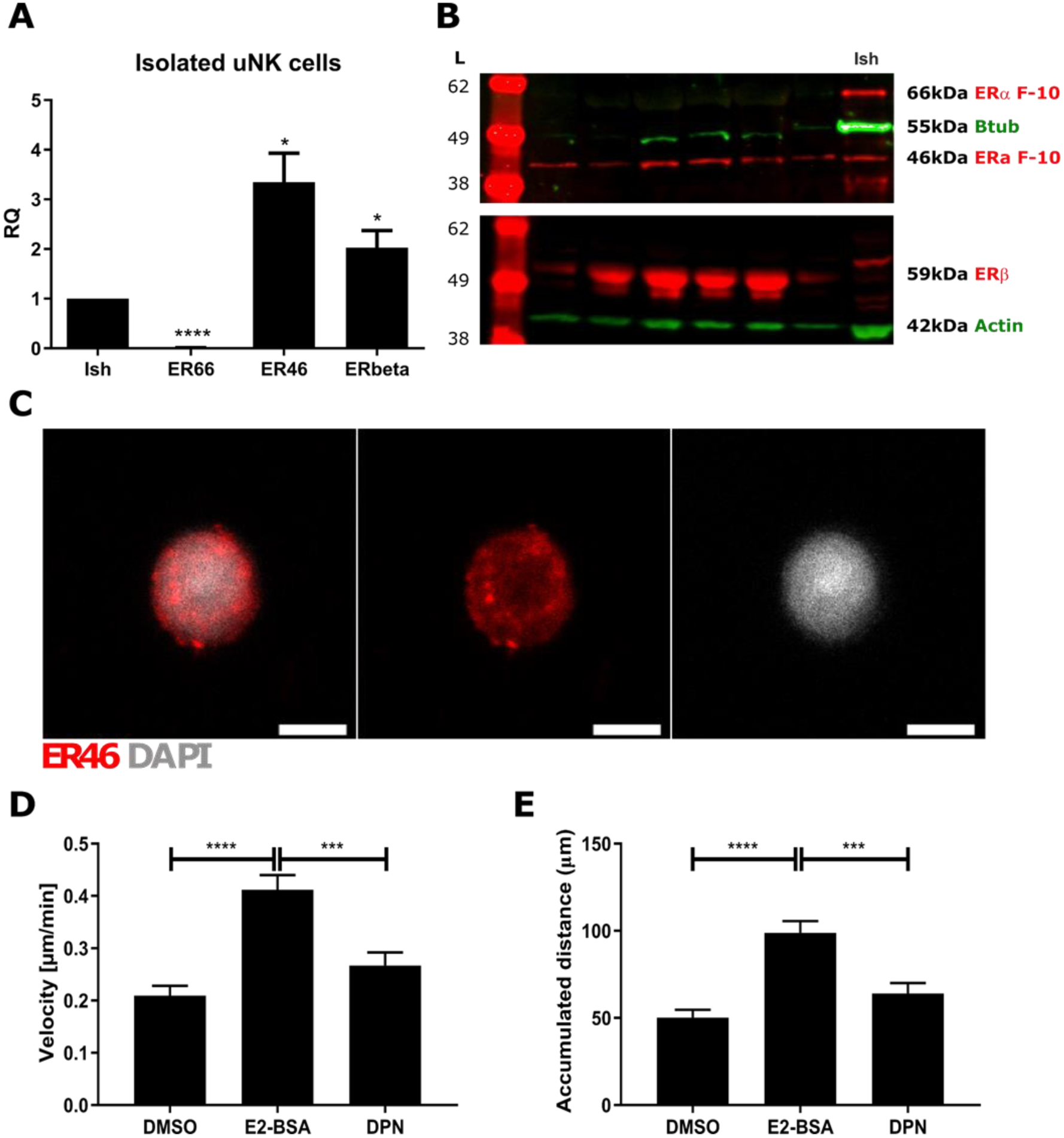
Isolated uNK cells express ER46 and increase cell motility in response to E2-BSA. UNK cells were isolated from decidua tissues by magnetic cell sorting using the MACS system. The expression of ER isoforms was assessed by qPCR, western blot and immunofluorescence. **A** Primers that mapped to either the N- or C-terminal of *ESR1*, or *ESR2* were used to assess mRNA expression in uNK cells relative to Ishikawa cell control lysates. The expression of mRNAs encoding the N-terminal of *ESR1* were significantly reduced in uNK cells compared to Ishikawa control (p<0.0001). In contrast, the expression of mRNAs encoding the C-terminal of *ESR1* and *ESR2* were significantly increased in uNK cells compared to Ishikawa control (p<0.01). **B** Protein expression was assessed by western blot in cell lysates of isolated uNK cells. ERα protein was assessed using the F-10 ERα antibody and a 46kDa band was detected in uNK cells. No corresponding 66kDa band was detected in uNK cells but was present in Ishikawa control lysate (Ish). ERβ was detected in both uNK cells and Ishikawa control by western blot. **C** Direct immunofluorescence was performed on isolated uNK cells using the F-10 ERα antibody and expression of ER46 was confirmed. Live cell imaging of isolated uNK cells was performed to assess cell motility in response to either vehicle control (DMSO), a membrane impermeable form of E2 (E2-BSA) or the ERβ selective agonist DPN. **D** E2-BSA significantly increased uNK cell velocity compared to vehicle control (DMSO; p<0.0001) and DPN (p<0.001). **E** E2-BSA significantly increased the accumulated distance of uNK cells compared to vehicle control (DMSO; p<0.0001) and DPN (p<0.001). DPN did not have an independent impact on either velocity or accumulated distance of uNK cells. ER antibodies and band sizes as indicated, loading controls B-actin or B-tubulin as indicated. Scale bars 5 µm, nuclear counterstain DAPI (grey). \**P* < 0.05, \*\*\**P* < 0.001, \*\*\*\**P* < 0.0001. Samples and analysis - qPCR: uNK cells, *n* = 5; Ishikawa *n* = 7; One-sample *t* test with hypothetical mean of 1. Western blot: uNK cells, *n* = 6. Cell motility analysis: *n* = 76 per treatment, Kruskal–Wallis test with multiple comparisons. All data are presented as mean ± s.e.m.

The human *ESR1* gene exhibits differential promoter usage and alternative splicing which give rise to splice variant isoforms of the receptor protein. ER46 was the first identified splice variant of human *ESR1* (initially designated hERα-46; (Flouriot, et al., 2002)). The ER46 variant is a 46kDa protein which lacks the N-terminal 173 amino acids of the full length ERα protein (66KDa, hereafter referred to as ER66) and arises from splicing of exon 1E to exon 2 via the E and F promoters (Flouriot, et al., 2002). ER46 contains both ligand binding and DNA binding domains and has been reported to bind oestradiol (E2) and to induce expression of oestrogen response element (ERE)-driven reporter genes (Flouriot, et al., 2002). ER46 and ER66 share identical sequence homology except that the N-terminal 173 amino acids of ER66 are absent in ER46. As all amino acids in ER46 are also present in ER66, there is no specific antibody that can uniquely identify ER46. It is therefore challenging to assess cell-specific patterns of native ER46 protein expression and this has limited our understanding of its functional significance.

ER46 and ER66 proteins can be resolved by size using western blotting techniques in combination with ERα-specific antibodies that recognise epitopes in the N-terminus (ER66 alone) or C-terminus (ER66 and/or ER46) of the proteins. Using this approach native expression of ER46 has been reported in human endothelial cell lines (Li, et al., 2003) and in human peripheral blood leukocytes (Pierdominici, et al., 2010). Detailed microscopy studies by Kim et al, using fluorescent tagged fusion protein revealed that ER46 can be detected localised to the plasma membrane in endothelial cells (Kim, et al., 2011). These studies have also demonstrated that membrane-associated ER46 can mediate rapid responses to oestrogens suggesting this receptor isoform may play a key role in ‘non-genomic’ or ‘membrane-initiated’ oestrogen receptor signalling (Kim, et al., 2014).

Uterine natural killer (uNK) cells are an abundant leukocyte population present in the endometrium during the late secretory phase and in the decidua of pregnancy and are characterised by high expression of the glycoprotein neural cell adhesion molecule (CD56) (Bulmer, et al., 1991, Koopman, et al., 2003). They are abundant in perivascular and luminal regions of the endometrium and play key roles in regulating vascular remodelling in early pregnancy and during placentation (Bulmer, et al., 2012, Robson, et al., 2012). Dysregulation of uNK cell function has been implicated in disorders of pregnancy including pre-eclampsia, foetal growth restriction, and recurrent pregnancy loss (Bulmer, et al., 2019, Gaynor, et al., 2017). However, the factors that regulate uNK cell function in both normal and pathological pregnancy remain poorly understood.

We have previously shown that isolated human uNK cells are exquisitely sensitive to oestrogens and can be stimulated to increase cell motility (chemokinesis and migration) in response to E2 (Gibson, et al., 2015). Notably, changes in uNK cell motility in response to E2 are rapid, initiated within minutes, and detected within 1 hour of treatment; consistent with a possible non-genomic signalling response (Gibson, et al., 2015). We demonstrated that uNK cell response to E2 was abrogated in the presence of the ER antagonist ICI 182,780 (Fulvestrant) consistent with an ER-dependent mechanism (Gibson, et al., 2015). We have previously characterised human uNK cells as ERα (ER66)-negative and ERβ-positive by immunohistochemistry and qPCR (Gibson, et al., 2015, Henderson, et al., 2003) but in those studies we did not consider expression of ER46 or its potential role in rapid responses to oestrogens.

In the current study we used qPCR, western blot and multiplex immunohistochemistry to assess expression of ER46, ER66 and ERβ in endometrial tissues and isolated uNK cells. We sought to identify cell populations within the endometrium that express ER46, define cellular localisation of receptor proteins and to investigate a potential functional role for ER46 in mediating oestrogen responses in uNK cells.

## Material and Methods

### Human Tissue Samples

Human endometrial tissues were obtained from women undergoing surgery for benign gynaecological conditions (n=24) and human decidua samples from women undergoing surgical termination of pregnancy, mean gestation of 10 weeks, (n=30). Local ethical committee approval was granted and written informed patient consent was obtained prior to tissue collection by a dedicated research nurse (Ethical approval held by HODC; LREC/05/51104/12 and LREC/10/51402/59). Tissue samples were fixed in 4% neutral buffered formalin or RNA Save (Geneflow, Staffordshire, UK). Stage of the menstrual cycle was determined histologically by an experienced gynaecological pathologist and by measurement of serum E2 and progesterone levels as previously detailed (Bombail, et al., 2008). Primary human uNK cells were isolated from fresh human first trimester decidua as described previously (Gibson, et al., 2015). Briefly, decidual tissues (n=18) were minced, digested in collagenase/DNAse and passed through 70 and 40 µm cell strainers. The cell suspension was overlaid on Histopaque 1077 (Sigma-Aldrich, USA) to separate leukocytes. UNK cells isolated by MACS magnetic bead separation using CD3 depletion and CD56 selection (Miltenyi Biotech, Germany).

Ishikawa (human endometrial adenocarcinoma) cells (ECACC_99040201) which express ER66 were used as a positive control for western blotting and qPCR: cells were cultured according to established protocols (Collins, et al., 2009).

### RNA Extraction, cDNA synthesis and Quantitative real time PCR

Total RNA was extracted from cell pellets or 20mg of tissue using Tri-Reagent and chloroform and homogenisation using a tissue lyser for 2 minutes at 20Hz (Qiagen). RNA was extracted using RNeasy Mini kit (Qiagen, UK) according to manufacturer’s instructions. RNA quantity and purity was confirmed by Nanodrop ND-1000 spectrophotometry (Thermo Scientific) and was standardised to 100ng/µl for all samples. CDNA was synthesised using SuperScript VILO cDNA Synthesis kit (Invitrogen).

Quantitative real time PCR (qPCR) was performed with primer sets (Supplemental Table 1) designed using Roche Universal Probe Library Assay Design Center (Roche Diagnostics, UK) in conjunction with corresponding FAM-labelled probes. Briefly, a reaction mix was prepared containing 1x Express Supermix, ribosomal 18S, 200nM of forward/reverse primer and 100nM probe. Samples were assayed in duplicate, using 18S as internal control reference gene on a 7900HT Fast Real Time PCR machine (Applied Biosytems). Amplification was performed at 95°C for 10 minutes then 40 cycles of 95°C for 15 seconds and 60°C for 1 minute. Target gene expression was assessed using the 2^-ΔΔCt^ method where the mean value of the proliferative samples (tissues) or Ishikawa cell homogenate (cell samples) was used for relative quantification.

### Protein Extraction

Total protein from 30mg of frozen tissue or cell pellets was extracted by homogenising in lysis buffer [1% Triton X-100, 167mM NaCl, 5mM EDTA (pH8.5), 50mM Tris (pH 7.5) 2µg/ml Aprotinin and 1x Halt protease inhibitor cocktail (Thermo Scientific)] using a Tissue Lyser for 2 minutes at 20Hz, followed by centrifugation at 13,000rpm (Eppendorf 5414R) for 10 minutes at 4°C. Ishikawa cell nuclear and cytoplasmic protein fractions were extracted using Nuclear Extraction Kit (Active Motif, Belgium) according to manufacturer’s instructions. Protein quantification was performed using DC protein Assay from Bio-Rad and read at 690nm on mass spectrophotometer (ThermoFisher, US).

### Western blot

Western blot was performed to identify ERα proteins corresponding to full length (66kDa) or truncated ERα (46kDa). Proteins were separated on NuPage Novex 4–12% Bis-Tris polyacrylamide gels (Life Technologies Inc.) under reducing conditions with NuPage MOPS SDS running buffer then transferred onto Immobilon FL transfer membrane (EMD Millipore) using a semidry blotter for 90 minutes at 14V. Membranes were incubated overnight at 4°C with primary antibodies: mouse anti-ERα 6F11 (1:300); mouse anti-ERα F-10 (1:1000); rabbit anti-ERβ (1:200); and loading controls were mouse anti-β-Tubulin (1:1000); mouse anti-β Actin (1:2000); rabbit anti-β Actin (1:500) respectively (Supplemental Table 2). Membranes were washed in PBS containing 0.1% Tween-20, incubated with appropriate species-specific fluorescent-conjugated secondary antibodies (Supplemental Table 3) and visualised using Licor Odyssey infrared imaging system (Licor).

### Immunohistochemistry

Tissues were sectioned and subjected to antigen retrieval in 0.01M citrate pH6 and immunohistochemistry performed according to standard methods (Critchley, et al., 2001). Sections were incubated overnight with primary antibodies; ERα (F-10), ERα (6F11) or CD56 (as detailed in Supplemental Table 2) at 4°C followed by incubation with peroxidase conjugated secondary antibody for 1 hour (Supplemental Table 3). Antigen detection was performed using Tyramide signal amplification (Perkin Elmer-TSA-Plus Fluorescein) according to manufacturer’s instruction. Negative controls, omitting the primary antibody, were included in each experiment.

For multiplex immunofluorescence experiments, an elution step was performed prior to addition of the next primary antibody by microwaving sections in 0.01M citrate buffer (pH 6.0) for 150 seconds and left to cool for 20min. This was followed by serum block and overnight incubation at 4°C with primary antibodies. Up to three primary antibodies, ERα (F-10), ERα (6F11) or CD56, were used and combined with PerkinElmer-TSA-Plus-Fluorescein (Green), PerkinElmer-TSA-Plus-Cy3 (Red) and PerkinElmer-TSA-Plus-Cy5 (Blue) respectively. Slides were counterstained with DAPI and mounted with Permafluor (Thermo Scientific) prior to imaging.

### Immunocytochemistry

Isolated uNK cells were cultured in coated BD Falcon Chamber slides (BD Bioscience, UK) and washed twice with PBS at room temperature. Cells were fixed in ice cold methanol for 20 minutes, washed, and permeabilised in a solution containing 0.2% IGEPAL (Sigma Aldrich, USA), 1% BSA and 10% NGS diluted in PBS for 20 minutes at room temperature. Endogenous peroxidase was blocked by immersing slides in 0.15% H_2_O_2_ in methanol for 30 minutes and non-specific binding was blocked by incubating cells in NGS/PBS/BSA for 30 minutes. Cells were incubated with an anti-ERα antibody (F-10) overnight followed by Cy3 Tyramide signal amplification (PerkinElmer-TSA-Plus-Cy3) according to manufacturer’s instructions. Slides were counterstained with DAPI and were mounted in Permafluor prior to imaging.

### Imaging

Fluorescent images were acquired with a Zeiss LSM 710 Confocal microscope and processed with ZEN 2009 Software (Zeiss).

### Live Cell Imaging

The chemokinesis of uNK cells was assessed as described previously (Gibson, et al., 2015). Isolated uNK cells were suspended in a collagen matrix in Ibidi μ-Slide Chemotaxis3D chamber slides (Ibidi, 80326, supplied by Thistle Scientific Ltd, Uddingston, UK). Chamber slides were set up containing serum-free phenol red-free RPMI 1640 media and treatment. The response to the membrane impermeable ligand E2-BSA (10nM equivalent), the ERβ-selective agonist 2,3-bis (4-hydroxyphenyl)-propionitrile (DPN; 10nM) or vehicle control (DMSO) was measured using time lapse microscopy. Cells were imaged every 2 minutes for 2 hours using Axiovert 200 Inverted Fluorescent Microscope (Zeiss). Data were analysed using ImageJ (manual cell tracking plug-in) and chemotaxis and migration tool software (Ibidi).

### Statistics

Statistical analysis was performed using GraphPad Prism. Kruskal-Wallis test with Dunn’s multiple comparison test was used to determine significance between treatments. Where data were analysed as fold change, significance was tested using one-sample *t* test with hypothetical mean of 1. Criterion for significance was p<0.05. All data are presented as mean ± SEM.

## Results

### Profiling human endometrial tissues reveals distinct patterns of ER isoform expression

Due to the overlapping sequence homology between mRNAs encoding full length ERα (ER66) and the truncated splice variant isoform ER46 it is impossible to design oligonucleotide primers that can uniquely distinguish between the two isoforms. In this study we designed primer pairs (see Supplementary Table 1) directed against sequences in the N or C-terminal of the receptor and used these to detect mRNAs for either ER66 alone (N-terminal primers) or ER66 and/or ER46 (C-terminal primers). Consistent with data from our previous studies (Critchley, et al., 2002, Milne, et al., 2005) and those of others (Binder, et al., 2015) mRNAs encoded by *ESR1* assessed using N-terminal primers were present in endometrial tissue homogenates from proliferative and secretory phase endometrium (Figure 1A) and significantly decreased in decidual tissue homogenates (p<0.01). In contrast, mRNA expression of *ESR1* assessed using C-terminal primers was detected in all samples and was most abundant in secretory phase endometrium (Figure 1B). Consistent with our previous findings (Critchley, et al., 2002), mRNAs encoded by *ESR2* (detected using primers directed against the wild type isoform, ERβ1) were more abundant in secretory phase endometrium and decidua than samples from proliferative phase (Figure 1C).

We next assessed protein expression of ER46 and ER66 isoforms by performing western blotting using antibodies directed against either the whole receptor (clone 6F11) or an epitope in the C-terminal domain (clone F-10) of ERα. Densitometry measurements confirmed variation in the abundance of ER proteins (Figure 2A and B); endometrial tissues expressed all three proteins whereas in decidua ER66 was not detected and expression of ER46 (p<0.05) and ERβ1 (p<0.001) was significantly greater than ER66. A single protein band (∼66kDa) was detected in endometrial tissue homogenates using the ERα 6F11 antibody: decidual tissue homogenates had no detectable protein at this size (Figure 2C and D). Using the C-terminal-specific ERα F-10 antibody, proteins corresponding to both 46kDa and 66kDa were detected in endometrium (Figure 2C) but only a 46kDa protein was detectable in decidua (Figure 2D). A single 59KDa band corresponding to full length ERβ1 (Critchley, et al., 2002) was detected in all samples (Figure 2C and D).

### Immunostaining of ER isoforms in endometrial tissues

Dual immunohistochemistry was performed in endometrial tissues to assess the pattern of expression of proteins recognised by the 6F11 and F-10 ERα antibodies. Expression of ER46 was inferred from the presence of staining using the C-terminal ERα F-10 antibody and absence of staining with the ERα 6F11 antibody. In proliferative phase endometrium (Figure 3), ERα was detected with both the ERα 6F11 antibody (green) and C-terminal ERα F-10 antibody (red). Consistent with our previous findings (Bombail, et al., 2008, Milne, et al., 2005), expression of ER66 was detected in nuclei of both stromal and epithelial cells (yellow arrows). In contrast, a divergent pattern of expression was observed in secretory phase tissue (Figure 3; ‘secretory’). ER66 detected using the ERα 6F11 antibody (green) was localised exclusively to cell nuclei and detected in all epithelial cells and some stromal cells. Positive staining using the C-terminal ERα F-10 antibody (ER46/66, red) was detected in the nuclei of epithelial and stromal cells and overlapped with ERα 6F11 antibody (yellow arrow). However, C-terminal ERα F-10 antibody (red) also localised to extra-nuclear sites and was detected in the cytoplasm of epithelial and stromal cells (white arrowhead) as well as the membrane of some cells (white arrows). Notably, when protein was localised to the membrane no staining was detectable in the nucleus using either antibody (white arrows). This pattern of expression was most obvious in first trimester decidual tissues which express lower concentrations of ER66 (Figure 1A and Figure 2B). Cytoplasmic expression of ERα was detected using the C-terminal ERα F-10 antibody (red) in decidualised stromal cells (white arrowhead) and membrane expression was apparent on numerous cells within the stromal compartment (white arrow).

### Human uNK cells express of ER46 in first trimester decidual tissues

As CD56-positive uterine natural killer (uNK) cells are the most abundant leukocyte in first trimester decidual tissues, we investigated whether membrane ERα expression was associated with this cell by performing triple immunohistochemistry using the 6F11 and ERα F-10 antibodies and anti-CD56 (Figure 4). Co-staining of ERα 6F11 antibody (green) confirmed our previous finding that CD56-positive cells (blue) were ER66-negative (Figure 4). In contrast, ERα was detected on the membranes of uNK cells using the C-terminal ERα F-10 antibody (red): this co-expression is visible as pink staining on the surface of uNK cells (white arrows; Figure 4).

### Expression of ER46 in isolated human uNK cells

UNK cells were isolated from decidua by magnetic sorting and expression of ER66, ER46 and ERβ1 was assessed by qPCR, western blot and immunocytochemistry (Figure 5). Consistent with our previous studies, mRNAs detected using N-terminal primers (ER66) were significantly lower in uNK cells than Ishikawa cells (p<0.0001), in contrast, mRNAs detected using C-terminal-specific primers were significantly higher in uNK cells (Figure 5A; p<0.05). Expression of ER46 in isolated uNK cells was confirmed by western blot (Figure 5B) and immunofluorescence with staining localised to the cell membrane (Figure 5C). Consistent with our previous findings, uNK cells contained mRNAs encoded by *ESR2* as well as protein of 59KDa on western blots corresponding to full length ERβ1 protein (Figure 5A and B).

### ER46 expression in uNK cells promotes membrane-initiated changes in cell motility

We have previously demonstrated that treatment of isolated uNK cells with E2 results in increased rates of cell migration (Gibson, et al., 2015). Based on receptor expression profiling described above, we investigated whether the impact of E2 on uNK cells could be mediated by ER46 (membrane) or ERβ (nucleus). Cells were treated with E2 conjugated to BSA (E2-BSA) which cannot cross the cell membrane and would putatively activate ER46, the ERβ-selective agonist DPN or to vehicle control (DMSO). We performed live cell imaging of uNK cells using time-lapse microscopy and assessed cell motility. E2-BSA significantly increased uNK cell velocity compared to both DMSO (p<0.0001) and DPN (p<0.001) (Figure 5D). Mean velocity of DPN-treated cells was slightly greater than DMSO, but this was not statistically significant. Similarly, E2-BSA significantly increased the accumulated distance of uNK cells compared to both DMSO (p<0.000 1) and DPN (p<0.001) (Figure 5E). DPN did not have an independent effect on uNK cell accumulated distance within 2 hours of incubation.

## Discussion

Oestrogens are essential regulators of endometrial function and fertility. Expression of the ER splice variant ER46 has been demonstrated in peripheral blood leukocytes and isolated endothelial cells. In the current study, we have shown for the first time that expression of ER46 in human endometrium is distinct from that of full length ERα (ER66). Notably, ER46 is uniquely expressed on the membrane of uNK cells which are otherwise ER66-negative. Functional analysis of uNK cells demonstrated that targeting of ER46 with E2-BSA increased cell motility via rapid, putatively non-genomic mechanisms.

Expression of ER46 has not previously been described in human endometrial tissues however by using antibodies able to distinguish between this variant and full length ER66 we identified ER46 protein in tissue homogenates from both cycling (non-pregnant) endometrium as well as first trimester decidua. Previous studies have reported that ER46 acts as a dominant negative repressor of ER66, inhibiting E2-induced transcription of a reporter gene and cell proliferation (Li, et al., 2003, Penot, et al., 2005). We suggest expression of ER46 is most likely to impact on classical responses to ER ligands in the endometrium during the proliferative phase when ER46 and ER66 were both detected in the nuclei of endometrial cells (Figure 3). In secretory phase and decidua tissue samples, immunohistochemistry demonstrated that ER46 was most abundant in the cytoplasm, whereas ER66 was exclusively nuclear. Thus, fewer positive cells were detected in which both ER46 and ER66 isoforms were co-localised to cell nuclei. This change in cellular localisation of ER46 indicates ER46/ER66 interactions may differ across the menstrual cycle. Further studies are needed to assess whether ER46/ER66 dimerisation can impact on the regulation endometrial function in response to E2. This may be particularly relevant to the pathophysiology of endometrial hyperplasia/cancer where oestrogens are key drivers of epithelial proliferation and cancer growth (Collins, et al., 2009, Sanderson, et al., 2017).

UNK cells are the predominant leukocyte in secretory endometrium and first trimester decidua where they mature to acquire phenotypic properties that distinguish them from their peripheral blood (pb) NK cell precursors (King, et al., 1991, King, et al., 1989, Pace, et al., 1989). Notably, uNK cells are transcriptionally distinct from pbNK cells (Koopman, et al., 2003) and they exhibit decreased cytotoxicity and increased cytokine secretion compared to pbNK cell subsets. This phenotype is critical to their function and they are essential mediators of vascular remodelling in early pregnancy (Robson, et al., 2012). Accumulating evidence supports a role for oestrogens in controlling the function of both pbNK precursors and their uNK cell descendants within the endometrium. Human pbNK cells are ER-positive with evidence for ERα and ERβ expression (Pierdominici, et al., 2010). Profiling of human pbNK cells isolated from different phases of the menstrual cycle demonstrated that pbNK cells exhibit increased adhesion on day 14 (when E2 concentrations peak in the circulation) compared to other phases of the cycle. In the same study, E2 treatment in vitro increased adhesion of pbNK cells to uterine tissues sections (van den Heuvel, et al., 2005). NK cells play a crucial role in defence against pathogens by carrying out cell-mediated toxicity. This function also appears to be regulated by oestrogens in women as NK cell activity, measured by lytic effector function, is reported to be increased in postmenopausal women (low circulating E2) compared to premenopausal women. Furthermore, NK cell activity is decreased in postmenopausal women following oestrogen hormone replacement therapy (Albrecht, et al., 1996). This effect is also mirrored in mouse splenic NK cells where E2 is reported to decrease cytotoxic activity (Curran, et al., 2001) and their proliferative capacity (Hao, et al., 2008). Thus, bioavailability of E2 appears to have impacts on homing of pbNK cells to the uterus and also to promote a low cytotoxicity phenotype that is similar to uNK cells. The oestrogen-dominated microenvironment found in the endometrium in early pregnancy is therefore likely to contribute to a similar functional adaptation of uNK cells within the tissue (Gibson, et al., 2013).

We previously demonstrated that incubation with E2 increases uNK cell motility (Gibson, et al., 2015), attributing the impact of E2 to signalling via ERβ1 as we failed to detect any ERα (ER66) in these cells. However, the changes in uNK cell motility detected in response to E2 were rapid (within 1 hour in (Gibson, et al., 2015)) which prompted us to consider a role for ER46 and membrane-initiated signalling as a mechanism to explain these changes in uNK cell function. Cells which express ER46 may be more likely to transduce oestrogenic responses via cell membrane-initiated pathways. For example, in ER-negative COS7 cells, in which expression of either ER46 or ER66 was induced, ER46 was found to be less efficient at inducing transcription of an ERE-reporter construct than ER66 but *more* efficient at inducing membrane-initiated phosphorylation of eNOS (Li, et al., 2003). ER46 has been located to the cell membrane of peripheral blood leukocytes and is reported to be the only ER isoform detected in membrane of pbNK cells (Pierdominici, et al., 2010). Stimulation of activated human pbNK cells with E2-BSA is reported to increase secretion of interferon-γ (Pierdominici, et al., 2010). In the current study we have detected ER46 protein on the cell membrane of decidual uNK cells and found that incubation with E2-BSA, but not the ERβ1-selective agonist DPN, rapidly increased cell motility consistent with ER46 mediating membrane-initiated rapid responses to oestrogens. We have previously demonstrated that oestrone (E1) and E2 are secreted by decidualised stromal cells which may account for accumulation of uNK cells in perivascular areas of the endometrium (Gibson, et al., 2013) where they promote vascular remodelling in early pregnancy (Gibson, et al., 2015, Robson, et al., 2012). Changes in uNK cell motility via ER46 may therefore be required for appropriate control of spatiotemporal remodelling during the establishment of pregnancy.

It is possible that expression of other ERs such as ER36 (36KDa ER isoform) or G protein-coupled ER (GPER) may mediate membrane-initiated responses to oestrogens in endometrial cells as both receptors have been detected at the cell membrane (Thomas, et al., 2005, Wang, et al., 2005, Wang, et al., 2006). However, to the best of our knowledge neither receptor has been detected in uNK cells. Furthermore, ER36 expression has not been reported in the endometrium and the receptor protein lacks both transcriptional activation domains (AF-1 and AF-2) (Wang, et al., 2005) and cannot bind E2 (Lin, et al., 2013). Whilst GPER has been detected in endometrial tissues its expression appears higher in proliferative than secretory phase or decidua and it has been localised to epithelial cells (Kolkova, et al., 2010, Plante, et al., 2012). This pattern contrasts with the abundant expression of ER46 in multiple cell types in endometrium and decidua reported in the current study. Although GPER binds E2 it does not bind other endogenous oestrogens such as E1 or oestriol (E3) (Thomas, et al., 2005) and drugs that inhibit activation of nuclear ERs, including ICI 182,780, function as full agonists to GPER (Prossnitz, et al., 2011). Given the results in our previous studies demonstrating that *both* E1 and E2 increase uNK cell migration and that these effects are abrogated by ICI 182,780 it is unlikely that GPER is responsible for this rapid change in cell function.

### Conclusion

In the present study we provide new evidence for expression of human ER46 in the endometrium and decidua and highlight a role for this isoform in oestrogenic regulation of uNK cell function. Given the importance of uNK cells to regulating vascular remodelling in early pregnancy and the potential for selective targeting of ER46, this may be an attractive future therapeutic target in the treatment of reproductive disorders.

## Supporting information

Supplementary data: Table1-3; uncropped western blots

## Authors Roles

DAG designed and carried out experimental work and wrote the manuscript. CB-S carried out experimental work. AE-Z and FC carried out experimental work and wrote the manuscript. HODC provided clinical samples and revised the manuscript. PTKS designed the work, wrote and revised the manuscript.

Research costs and salaries (DAG, FC, AE-Z, and PTKS) were supported by MRC Programme Grants G1100356/1 and MR/N024524/1 to PTKS. HODC was supported by MRC grant G1002033.

## Acknowledgements

We thank members of PTKS group for support and technical assistance. We thank Research Nurses Catherine Murray and Sharon McPherson for patient recruitment and collection of tissues. We are grateful to Prof Alistair Williams for histological staging of endometrial tissues.

## References

Albrecht AE, Hartmann BW, Scholten C, Huber JC, Kalinowska W, Zielinski CC. Effect of estrogen replacement therapy on natural killer cell activity in postmenopausal women. Maturitas 1996;25: 217–222.

Binder AK, Winuthayanon W, Hewitt SC, Couse JF, Korach KS. Chapter 25 - Steroid Receptors in the Uterus and Ovary. In Plant TM and Zeleznik AJ (eds) Knobil and Neill’s Physiology of Reproduction (Fourth Edition). 2015. Academic Press, San Diego, pp. 1099–1193.

Bombail V, MacPherson S, Critchley HO, Saunders PT. Estrogen receptor related beta is expressed in human endometrium throughout the normal menstrual cycle. Human reproduction 2008;23: 2782–2790.

Bulmer JN, Innes BA, Levey J, Robson SC, Lash GE. The role of vascular smooth muscle cell apoptosis and migration during uterine spiral artery remodeling in normal human pregnancy. FASEB J 2012;26: 2975–2985.

Bulmer JN, Lash GE. Uterine natural killer cells: Time for a re-appraisal? F1000Research 2019;8.

Bulmer JN, Morrison L, Longfellow M, Ritson A, Pace D. Granulated lymphocytes in human endometrium: histochemical and immunohistochemical studies. Human reproduction 1991;6: 791–798.

Chantalat E, Boudou F, Laurell H, Palierne G, Houtman R, Melchers D, Rochaix P, Filleron T, Stella A, Burlet-Schiltz O et al. The AF-1-deficient estrogen receptor ERalpha46 isoform is frequently expressed in human breast tumors. Breast cancer research: BCR 2016;18: 123.

Collins F, MacPherson S, Brown P, Bombail V, Williams AR, Anderson RA, Jabbour HN, Saunders PT. Expression of oestrogen receptors, ERalpha, ERbeta, and ERbeta variants, in endometrial cancers and evidence that prostaglandin F may play a role in regulating expression of ERalpha. BMC Cancer 2009;9: 330.

Critchley HO, Brenner RM, Henderson TA, Williams K, Nayak NR, Slayden OD, Millar MR, Saunders PT. Estrogen receptor beta, but not estrogen receptor alpha, is present in the vascular endothelium of the human and nonhuman primate endometrium. The Journal of clinical endocrinology and metabolism 2001;86: 1370–1378.

Critchley HO, Henderson TA, Kelly RW, Scobie GS, Evans LR, Groome NP, Saunders PT. Wild-type estrogen receptor (ERbeta1) and the splice variant (ERbetacx/beta2) are both expressed within the human endometrium throughout the normal menstrual cycle. The Journal of clinical endocrinology and metabolism 2002;87: 5265–5273.

Curran EM, Berghaus LJ, Vernetti NJ, Saporita AJ, Lubahn DB, Estes DM. Natural killer cells express estrogen receptor-alpha and estrogen receptor-beta and can respond to estrogen via a non-estrogen receptor-alpha-mediated pathway. Cellular immunology 2001;214: 12–20.

Flouriot G, Brand H, Seraphin B, Gannon F. Natural trans-spliced mRNAs are generated from the human estrogen receptor-alpha (hER alpha) gene. The Journal of biological chemistry 2002;277: 26244–26251.

Gaynor LM, Colucci F. Uterine Natural Killer Cells: Functional Distinctions and Influence on Pregnancy in Humans and Mice. Frontiers in immunology 2017;8: 467.

Gibson DA, Greaves E, Critchley HO, Saunders PT. Estrogen-dependent regulation of human uterine natural killer cells promotes vascular remodelling via secretion of CCL2. Human reproduction 2015;30: 1290–1301.

Gibson DA, McInnes KJ, Critchley HO, Saunders PT. Endometrial Intracrinology--generation of an estrogen-dominated microenvironment in the secretory phase of women. The Journal of clinical endocrinology and metabolism 2013;98: E1802–1806.

Gibson DA, Saunders PT. Estrogen dependent signaling in reproductive tissues - a role for estrogen receptors and estrogen related receptors. Mol Cell Endocrinol 2012;348: 361–372.

Gibson DA, Simitsidellis I, Collins F, Saunders PTK. Endometrial Intracrinology: Oestrogens, Androgens and Endometrial Disorders. International journal of molecular sciences 2018;19.

Hao S, Li P, Zhao J, Hu Y, Hou Y. 17beta-estradiol suppresses cytotoxicity and proliferative capacity of murine splenic NK1.1+ cells. Cell Mol Immunol 2008;5: 357–364.

Henderson TA, Saunders PT, Moffett-King A, Groome NP, Critchley HO. Steroid receptor expression in uterine natural killer cells. The Journal of clinical endocrinology and metabolism 2003;88: 440–449.

Kim KH, Toomre D, Bender JR. Splice isoform estrogen receptors as integral transmembrane proteins. Molecular biology of the cell 2011;22: 4415–4423.

Kim KH, Young BD, Bender JR. Endothelial estrogen receptor isoforms and cardiovascular disease. Mol Cell Endocrinol 2014;389: 65–70.

King A, Balendran N, Wooding P, Carter NP, Loke YW. CD3-leukocytes present in the human uterus during early placentation: phenotypic and morphologic characterization of the CD56++ population. Developmental immunology 1991;1: 169–190.

King A, Wellings V, Gardner L, Loke YW. Immunocytochemical characterization of the unusual large granular lymphocytes in human endometrium throughout the menstrual cycle. Human immunology 1989;24: 195–205.

Kolkova Z, Noskova V, Ehinger A, Hansson S, Casslen B. G protein-coupled estrogen receptor 1 (GPER, GPR 30) in normal human endometrium and early pregnancy decidua. Molecular human reproduction 2010;16: 743–751.

Koopman LA, Kopcow HD, Rybalov B, Boyson JE, Orange JS, Schatz F, Masch R, Lockwood CJ, Schachter AD, Park PJ et al. Human decidual natural killer cells are a unique NK cell subset with immunomodulatory potential. J Exp Med 2003;198: 1201–1212.

Li L, Haynes MP, Bender JR. Plasma membrane localization and function of the estrogen receptor alpha variant (ER46) in human endothelial cells. Proceedings of the National Academy of Sciences of the United States of America 2003;100: 4807–4812.

Lin AH, Li RW, Ho EY, Leung GP, Leung SW, Vanhoutte PM, Man RY. Differential ligand binding affinities of human estrogen receptor-alpha isoforms. PloS one 2013;8: e63199.

Milne SA, Henderson TA, Kelly RW, Saunders PT, Baird DT, Critchley HO. Leukocyte populations and steroid receptor expression in human first-trimester decidua; regulation by antiprogestin and prostaglandin E analog. The Journal of clinical endocrinology and metabolism 2005;90: 4315–4321.

Pace D, Morrison L, Bulmer JN. Proliferative activity in endometrial stromal granulocytes throughout menstrual cycle and early pregnancy. Journal of clinical pathology 1989;42: 35–39.

Penot G, Le Peron C, Merot Y, Grimaud-Fanouillere E, Ferriere F, Boujrad N, Kah O, Saligaut C, Ducouret B, Metivier R et al. The human estrogen receptor-alpha isoform hERalpha46 antagonizes the proliferative influence of hERalpha66 in MCF7 breast cancer cells. Endocrinology 2005;146: 5474–5484.

Pierdominici M, Maselli A, Colasanti T, Giammarioli AM, Delunardo F, Vacirca D, Sanchez M, Giovannetti A, Malorni W, Ortona E. Estrogen receptor profiles in human peripheral blood lymphocytes. Immunology letters 2010;132: 79–85.

Plante BJ, Lessey BA, Taylor RN, Wang W, Bagchi MK, Yuan L, Scotchie J, Fritz MA, Young SL. G protein-coupled estrogen receptor (GPER) expression in normal and abnormal endometrium. Reproductive sciences 2012;19: 684–693.

Prossnitz ER, Barton M. The G-protein-coupled estrogen receptor GPER in health and disease. Nature reviews Endocrinology 2011;7: 715–726.

Robson A, Harris LK, Innes BA, Lash GE, Aljunaidy MM, Aplin JD, Baker PN, Robson SC, Bulmer JN. Uterine natural killer cells initiate spiral artery remodeling in human pregnancy. FASEB J 2012;26: 4876–4885.

Sanderson PA, Critchley HO, Williams AR, Arends MJ, Saunders PT. New concepts for an old problem: the diagnosis of endometrial hyperplasia. Human reproduction update 2017;23: 232–254.

Thomas P, Pang Y, Filardo EJ, Dong J. Identity of an estrogen membrane receptor coupled to a G protein in human breast cancer cells. Endocrinology 2005;146: 624–632.

van den Heuvel MJ, Horrocks J, Bashar S, Taylor S, Burke S, Hatta K, Lewis JE, Croy BA. Menstrual cycle hormones induce changes in functional interactions between lymphocytes and decidual vascular endothelial cells. The Journal of clinical endocrinology and metabolism 2005;90: 2835–2842.

Wang Z, Zhang X, Shen P, Loggie BW, Chang Y, Deuel TF. Identification, cloning, and expression of human estrogen receptor-alpha36, a novel variant of human estrogen receptor-alpha66. Biochemical and biophysical research communications 2005;336: 1023–1027.

Wang Z, Zhang X, Shen P, Loggie BW, Chang Y, Deuel TF. A variant of estrogen receptor-{alpha}, hER-{alpha}36: transduction of estrogen-and antiestrogen-dependent membrane-initiated mitogenic signaling. Proceedings of the National Academy of Sciences of the United States of America 2006;103: 9063–9068.

