## Supplementary data: Table1-3; uncropped western blots for "Profiling the expression and function of ER46 in human endometrial tissues and uterine NK cells"

Running title: ER46 in human endometrium

Supplementary data: Table 1-3; uncropped western blots

**Supplemental Table 1: Table of oligonucleotide sequences used in qPCR analysis**

| Primer Name | Accession number | Sequence | Primer position | UPL Probe |
|---|---|---|---|---|
| *ESR1* (N-terminal) Forward | Nm_000125 | *AACCAGTGCAACCATTGA TAAAA* | 1035-1056 | 69 |
| *ESR1* (N-terminal) Reverse | Nm_000125 | *TCCTCTTCGGTCTTTTCG TATC* | 1124-1145 | 69 |
| *ESR1* (C-terminal) Forward | Nm_0011227 41.1 | *TCTGGAAAGACGTTCTTG ATCC* | 121-146 | 29 |
| *ESR1* (C-terminal) Reverse | Nm_0011227 41.1 | *GGAGGGTCATGGTCATG GT* | 216-234 | 29 |
| *ESR2* Forward | Nm_001437 | *GCTCCTGTCCCACGTCA G* | 1848-1865 | 62 |
| *ESR2* Reverse | Nm_001437 | *TGGGCATTCAGCATCTCC* | 1944-1961 | 62 |

**Supplemental Table 2: Primary antibodies**

| Antibody name | Species | Supplier, Catalogue no. |
|---|---|---|
| ERα (6F11) | mouse | Vector, VP-614 |
| ERα (F-10) | mouse | Santa Cruz Biotechnology, sc-8002 |
| ERβ (H-150) | rabbit | Santa Cruz Biotechnology, sc-8974 |
| NCAM (CD56) | mouse | Invitrogen, 18-0152 |
| β-Actin | rabbit | Abcam, ab25894 |
| β-Actin | mouse | Sigma, A5441 |
| β-Tubulin (H-235) | rabbit | Santa Cruz Biotechnology, sc-9104 |
| β-Tubulin (AC-15) | mouse | Sigma-Aldrich, T4026 |

**Supplementary Table 2**: **Secondary antibodies**

| Antibody name | Species | Supplier, Catalogue no. | Dilution |
|---|---|---|---|
| Anti-rabbit biotinylated | goat | Vector Laboratories, BA-1000 | 1:500 |
| Anti-mouse IgG biotinylated | goat | Vector Laboratories, BA-9200 | 1:500 |
| Anti-mouse Peroxidase | goat | Dako, P0447 | 1:200 |
| IRDye 680 RD Anti-mouse | donkey | Licor, 926-68072 | 1:10000 |
| IRDye 680 RD Anti-rabbit | donkey | Licor, 926-68073 | 1:10000 |
| IRDye 800 CW Anti-mouse | donkey | Licor, 32212 | 1:10000 |
| IRDye 800 CW Anti-rabbit | donkey | Licor, 926-32213 | 1:10000 |
| Strepavidin (Horseradish peroxidase) | N/A | Vector Laboratories, SA-5004 | 1:500 |

### Supplementary western blot gel data

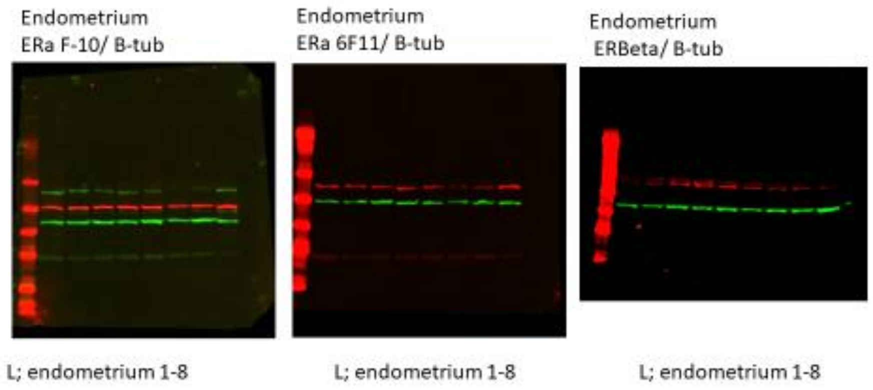


Figure 2, uncropped western blot gels

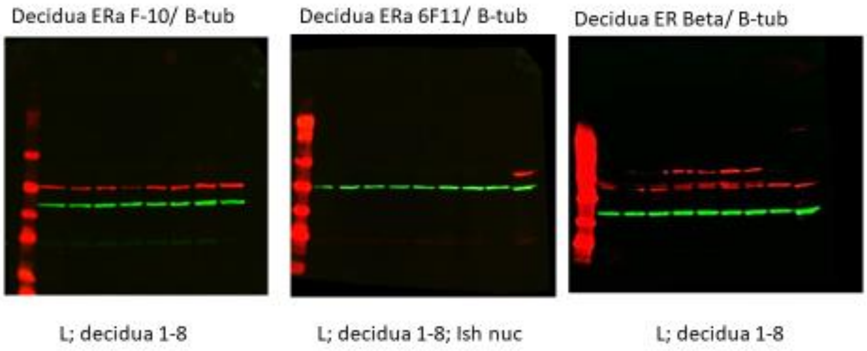


Figure 2, uncropped western blot gels

uNK ERa F-10/ B-tub | uNK ERa 6F11/ B-tub | uNK ER Beta/ B-tub

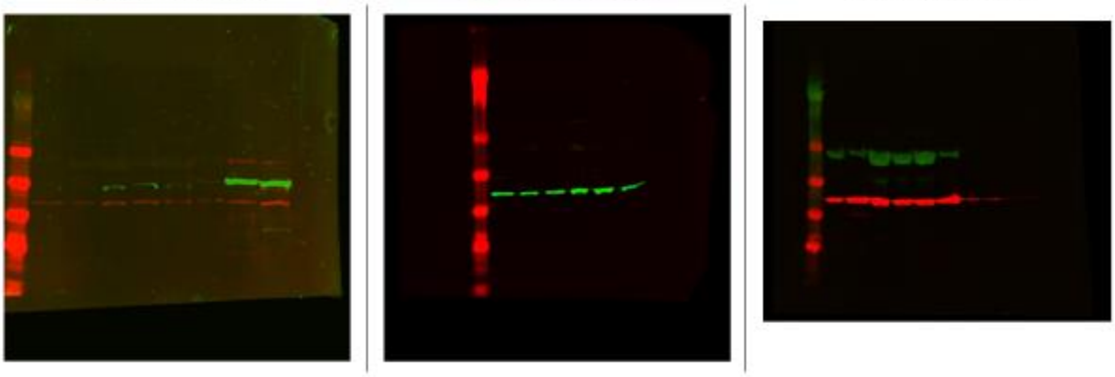

L; uNK1-6; Ish whole; Ish nuc | L; uNK1-6 | L; uNK 1-6

Figure 3, uncropped western blot gels
